# Sex-related gut microbiota in three geographically separated Norway lobster (*Nephrops norvegicus*) populations

**DOI:** 10.1101/2025.02.14.638244

**Authors:** Polina Rusanova, Eleni Nikouli, Michele Casini, Gioacchino Bono, Elena Mente, Alexandra Meziti, Konstantinos Kormas

## Abstract

Despite the ecological and economic interest of the Norway lobster (*Nephrops norvegicus*), its gut microbiota remains largely understudied. The current study aimed at investigating the gut bacterial microbiota in three geographically separated *N. norvegicus* populations from the Mediterranean and the North Sea and detecting any potential sex-related microbiota differences, by high-throughput sequencing of the V3-V4 16S rRNA gene diversity of the gut tissue. From the Greek population, egg-bearing females were also caught. A total of 2,385 operational taxonomic units (OTUs) were identified and between 417 and 1290 occurred in each population/sex group. The dominant OTUs belonged to the Fusobacteriia and Bacteroidia (Sweden), Bacilli and Gammaproteobacteria (Italy) and Spirochaetia and Bacilli (Greece) bacterial classes. In the eggs, the Actinobacteria, Alphaproteobacteria and Gammproteobacteria prevailed. Four OTUs related to the *Oceanispirochaeta*, *Kordiimonas, Desulfovibrio, Carboxylicivirga* genera and one unafilliated OTU were positively correlated (p values between 0.001 and 0.04) to body size indicating their potential role in the animal’s nutrition and growth. No statistically significant differences were found between males-females in each of the three populations. However, statistically significant differences between populations for each sex, were found for all females (p values between 0.008 and 0.032) and for the males between the most distant populations, i.e. Italy-Sweden (p=0.021) and Greece-Sweden (p=0.015). The eggs microbiota was statistically different from both the adult females (p=0.027) and males (p=0.046) gut microbiota. Overall, this study showed that *N. norvegicus* gut microbiota are differentiated between geographically distant populations and that sex-related differences are not significant.

## 1. INTRODUCTION

The hologenome and holobiont concepts (Zilber-Rosenberg and Rosenberg, 2008) have opened a new way we view animal life (Theis et al., 2016; Webster, 2017) for their ontogeny and function (Stencel and Wloch-Salamon, 2022; Troussellier et al., 2017). Thus, during the last few years, the scientific interest on holobiont research has boomed for animals, as well. However, there are yet animal species from various habitats, whose microbiomes remain largely unknow, despite their ecological and/or economic importance and the ongoing rapid technological advancements in DNA/RNA technologies.

One such species is the Norway lobster *Nephrops norvegicus* (Linnaeus, 1758) a decapod crustacean, langoustine or scampi, which is a typical clawed lobster with a slender body, long claws and large dark eyes. *Nephrops norvegicus* is considered one of the most important species for fisheries, and is considered a particularly valuable commercial crustacean species in Europe. It is a benthic crustacean dependent on muddy-type sediments suitable for the construction of burrows. It is widely distributed across the continental shelves of the northeast Atlantic Ocean and the Mediterranean Sea, from Iceland and Norway in the north, to Morocco and the Adriatic seas in the south (Bell et al., 2013). Populations dwelling in colder waters around Iceland and the Faroe Islands undertake a biennial breeding cycle whereas in the Mediterranean follow an annual breeding cycle (Bell et al., 2013). *N. norvegicus* are opportunistic predators and consume a wide variety of prey species including crustaceans, polychaetes, molluscs and echinoderms (Bell et al., 2013). *N. norvegicus* population genetics within the Mediterranean Sea using different methods revealed low or moderate genetic differentiation between geographical regions (Atlantic vs. Mediterranean) but no geographical pattern of genetic differentiation, thus genetic variability seems to be randomly distributed among populations (Maltagliati et al., 1998; Passamonti et al., 1997). Nevertheless, different populations were strongly related to spatial and environmental features of the Mediterranean (Commission et al., 2022).

The animal gut microbiome, i.e. the collective genetic material of all microbes found in the gut, is considered one of the central biological features for the reproduction, development, nutrition, growth, health/immunity of their host (Diwan et al., 2023; Singh et al., 2025) but even in the conservation of wildlife (Kanika et al., 2025; West et al., 2019). The microbiome concept is so established that the microbiome has been proposed as a means and a target of manipulation for improving farmed animals production (Luna et al., 2022). Gut microbiomes are also useful at assessing the impact of environmentally induced stress of animals (Evariste et al., 2019) but also as responders to such disturbances (See et al., 2025). Although decapods have attracted some scientific interest in regards to their microbiome (Foysal, 2023), even to date *N. norvegicus* as a holobiont has been sporadically studied. The bacterial microbiota of *N. norvegicus* has been investigated both in natural and experimentally reared populations (Meziti and Kormas, 2013; Meziti et al., 2012; Meziti et al., 2010) while pathogens such as the dinoflagellate *Hematodinium* (Small et al., 2006) have also been investigated. The above render the species’ microbiome largely unknown, especially in the framework of the most recently developed and high-throughput sequencing technologies. For these reasons, the current study aimed at (a) investigating the gut bacterial microbiota in three geographically separated *N. norvegicus* populations from the Mediterranean and the North Sea, (b) detecting any potential sex-related microbiota differences and (c) suggesting potential core bacterial taxa for the species populations. It is hypothesized that the animal’s gut bacterial communities are highly similar as the animals live in habitats with rather low seasonal variations and very similar environmental conditions and also to the little, if any, genetic differentiation between the populations.

## 2. MATERIALS AND METHODS

### 2.1. Sample collection

In the European Union, decapods are not classified as experimental animals under legislation (Directive 2010/63/EU and SFS 2019:66), therefore no animal ethical permits for their collection are required. In Sweden the samples for this study were collected during the ordinary International Bottom Trawl Survey, performed under the EU Data Collection Framework. In Italy the samples for this study were part of the daily landings of standard commercial fishing vessel operating in the Strait of Sicily therefore these samples are subject to IACUC regulations. In Greece the samples were acquired during onboard fishing operations with commercial bottom trawler in the frame of the National Data Collection Framework Program; by the time of capture all samples were already dead. All sampling efforts was limited to minimize large-scale impacts on the populations and is conducted using methodologies to ensure animal welfare and all speciments didn’t go through any experimental procedure.

Samples from the North Aegean Sea were collected southwest off Thasos Island (Greece, 40.46°N 24.6°E) between 190 and 435 m depth on 31 October 2023 by a bottom otter trawl boat. The Italian samples were collected in the Strait of Sicily (37° 16’ N, 13° 02’W, at 350-400 m depth) on 13 September 2023 by a commercial bottom trawler. Samples from the Kattegat, eastern North Sea (56.22°N 12.16°E, at 33 m depth), were collected by a research vessel equipped with a bottom trawling fishing net on 28 August 2023. From each sampling site, a total of seven male and seven females were collected for further analysis (Table S1, Figure S1).

### 2.2. Sample processing and microbiota analysis

For each individual and while on board the fishing vessel, the animals were placed on ice and the abdomen was severed from the cepaholothorax immediately after collection of the animals. The entire gut tissue was then carefully extracted by holding the abdomen with one hand and gently pulling the telson with the other. The collected gut tissue samples were stored immediately in DNA/RNA shield (Zymo Research, USA) and stored at -80^0^C after a few days. Gut bacterial communities composition of the gut tissue was determined after bulk DNA was extracted from approximately 0.25 mg of whole gut tissue (or eggs in the case for specimens from Greece) with the DNeasy PowerSoil Pro Kit (Qiagen, Germany) with no modifications from the suggested protocol. The V3–V4 regions of the bacterial 16S rRNA genes were amplified from the extracted DNA with the primer pair S-D-Bact-0341-b-S-17 and S-D-Bact-115 0785-a-A-21 (Klindworth et al., 2012). Sequencing of the amplicons was performed on a MiSeq Illumina instrument (2×300 bp) at the MRDNA Ltd. (Shallowater, TX, USA) sequencing facilities. The raw DNA sequences from this study have been submitted to the Sequence Read Archive (https://www.ncbi.nlm.nih.gov/sra/) in the BioProject PRJNA1070646 (BioSample SAMN39675726). The standard operating procedure of MOTHUR software (v.1.48.0) (Schloss et al., 2011; Schloss et al., 2009) was used for processing all the raw 16S rRNA sequence reads. Sequences assigned to mitochondria and chloroplasts, and single singletons were excluded from further analyses. The operational taxonomic units (OTUs) were determined at 97% cutoff similarity level and were classified with the SILVA database release 138.1 (Quast et al., 2013; Yilmaz et al., 2014). The final OTUs table was normalized to 33,164 sequence reads. Rarefaction curves (Figure S2) reached the plateau phase for the number of sequence reads we used for our analysis. The Nucleotide BLAST (http://blast.ncbi.nlm.nih.gov) tool was used for identifying the closest relatives of the resulting OTUs.

Testing of the differences across all samples were implemented in the PAleontological STudies (PAST) software (Hammer et al., 2001) by applying non-metric multidimensional scaling (nMDS) based on the unweighted pair group method with the arithmetic mean Bray–Curtis similarity. In addition, Bary-Curtis similarity permutational multivariate analysis of variance (PERMANOVA) with 9999 permutations, between the three geographical locations and sex of the *N. norvegicus* gut microbiota and eggs, was applied.

## 3. RESULTS

The three investigated populations had specimens of different sizes, based on their carapace length and body weight (Table S1). The smallest animals were caught in Italy and the largest in Greece. Statistically significant differences between females-males, based on PERMANOVA, occurred only for the animals from Sweden (p=0.027, F=19.810). Among animals of the same sex, the male specimens from Italy differed significantly from those from Sweden (p=0.011, F=125.100) while the females from Italy were different from those from Greece (p=0.015, F=32.600) and Sweden (p=0.014, F=277.100). The three populations were clearly separated with their male and female specimens overlapping in terms of their gut microbiota structure (Figure 1). The relative contributions of bacteria at the class-level were also similar between females and males in each of the three populations, with Fusobacteriia and Bacteroidia prevailing in the Swedish specimens, Bacilli and Gammaproteobacteria prevailing in the Italian specimens, and Spirochaetia along with Bacilli prevailing in the female Greece gut samples while the male samples were very diverse. In the eggs samples from the specimens from Greece, the Actinobacteria, Alphaproteobacteria and Gammproteobacteria prevailed (Figure 2).

**Figure 1.**
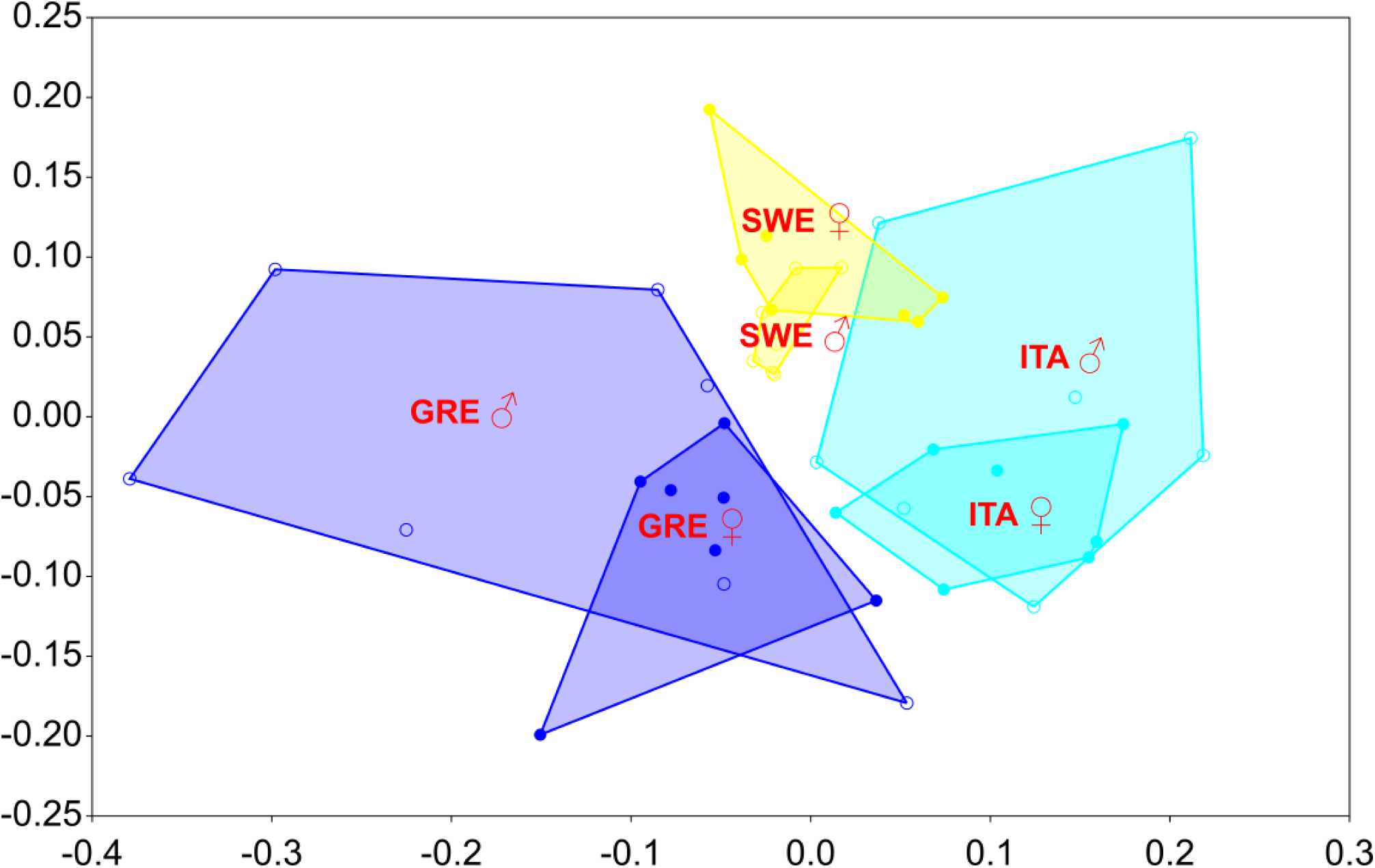
Non-metric multidimensional scaling (nMDS) of male (♂) and female (♀) individuals of *Nephrops norvegicus* gut from Greece (GRE), Italy (ITA) and Sweden (SWE).

**Figure 2.**
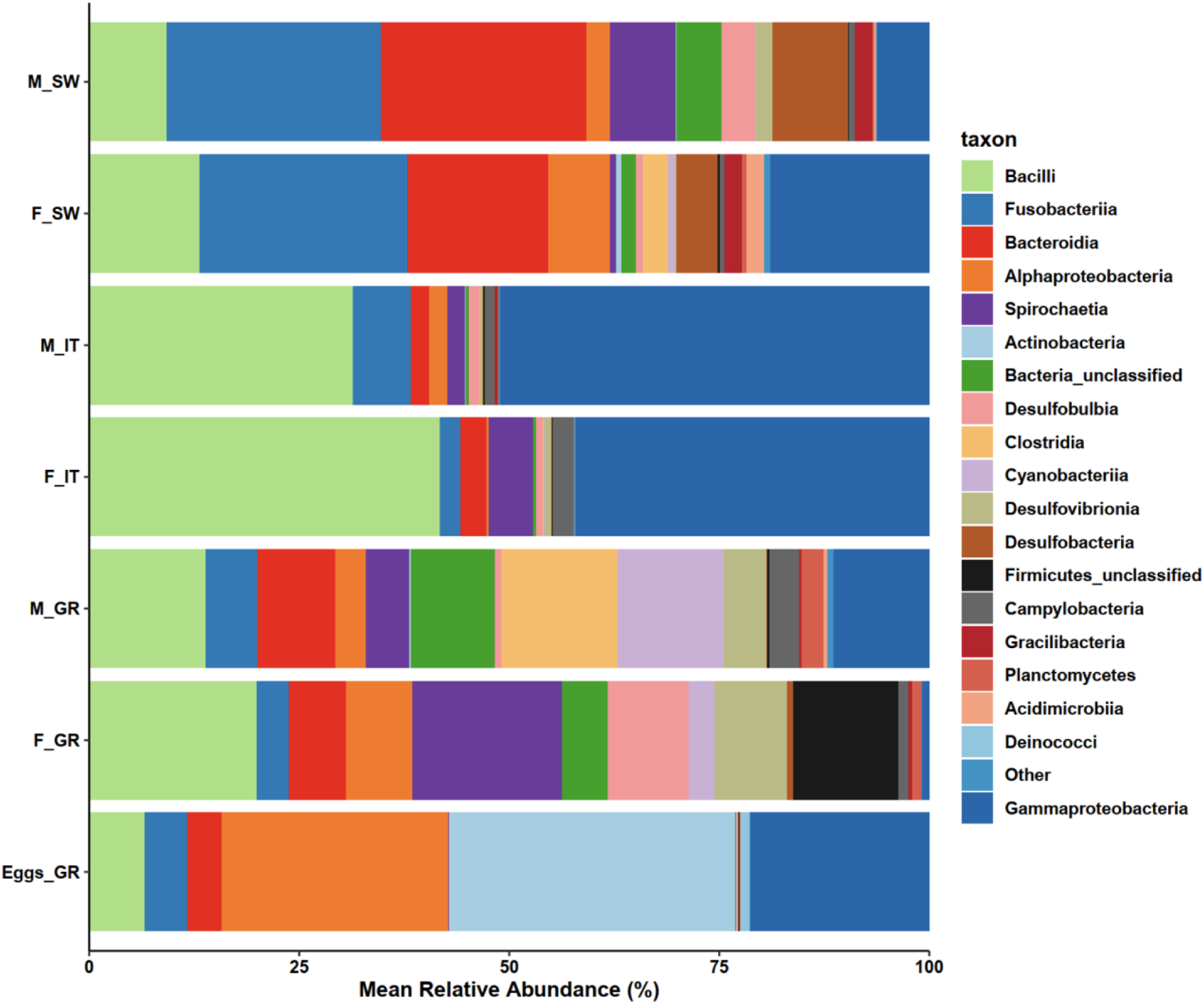
Relative abundance of class-level operational taxonomic units in the female (F) and male (M) *Nephrops norvegicus* gut from Greece (GR), Italy (IT) and Sweden (SW).

A total of 2,385 bacterial OTUs were identified across all samples. Females had between 417 (Greece) and 931 (Sweden) OTUs while the respective range for males was between 489 (Italy) and 1290 (Greece), while 180 OTUs were found in eggs from Greece (Table 1). The structure of the eggs bacterial communities was statistically different from both the gut microbiota of both females and males (Table 2). Moreover, there was no overlap between the dominant (relative abundance ≥80%) OTUs in these samples (Table S2). Females had higher number of shared OTUs with the eggs bacterial microbiota (24.9%) compared to males (10.6%) (Figure S3). For each population, no significant differences were observed between females and males, but statistically significant differences occurred for each sex between all populations except between males from Greece and Italy (Table 2).

**Table 1.** Midgut bacterial operational taxonomic units (OTUs) from three *Nephrops norvegicus* populations. Sequence reads=33,164; N=7 for male and female individuals, N=5 for eggs.

|  |  | Average carapace length<br>(mm) (coefficient of<br>variation) | Average total body<br>weight (g) (coefficient of<br>variation) | No. of OTUs<br>(average, coefficient of<br>variation) | No. of OTUs with<br>≥80% relative<br>abundance | Most dominant OTU<br>(% relative abundance) |
| --- | --- | --- | --- | --- | --- | --- |
| Greece | Eggs | - | - | 180<br>(86, 4.2%) | 17 | OTU-0020 (18.5%)<br>*Cutibacterium* |
|  | Females | 46.0 (11.6%) | 74.0 (30.9%) | 417<br>(128, 24.1%) | 11 | OTU-0004 (17.8%)<br>*Oceanispirochaeta* |
|  | Males | 46.7 (14.6%) | 77.0 (42.0%) | 1290<br>(335, 64.2%) | 20 | OTU-0018 (8.4%)<br>*Cyanobium* |
| Italy | Females | 34.9 (4.5%) | 32.4 (10.3%) | 434<br>(136, 24.0%) | 7 | OTU-0002 (20.2%)<br>*Photobacterium* |
|  | Males | 36.9 (4.0%) | 36.3 (13.9%) | 489<br>(153, 30.4%) | 10 | OTU-0002 (20.3%)<br>*Photobacterium* |
| Sweden | Females | 45.0 (0.0%) | 64.6 (2.5%) | 931<br>(252, 63.0%) | 14 | OTU-0001 (24.5%)<br>*Psychrilyobacter* |
|  | Male | 45.0 (0.0%) | 67.9 (1.6%) | 527<br>(161, 36.7%) | 11 | OTU-0001 (25.5%)<br>*Psychrilyobacter* |

**Table 2.**
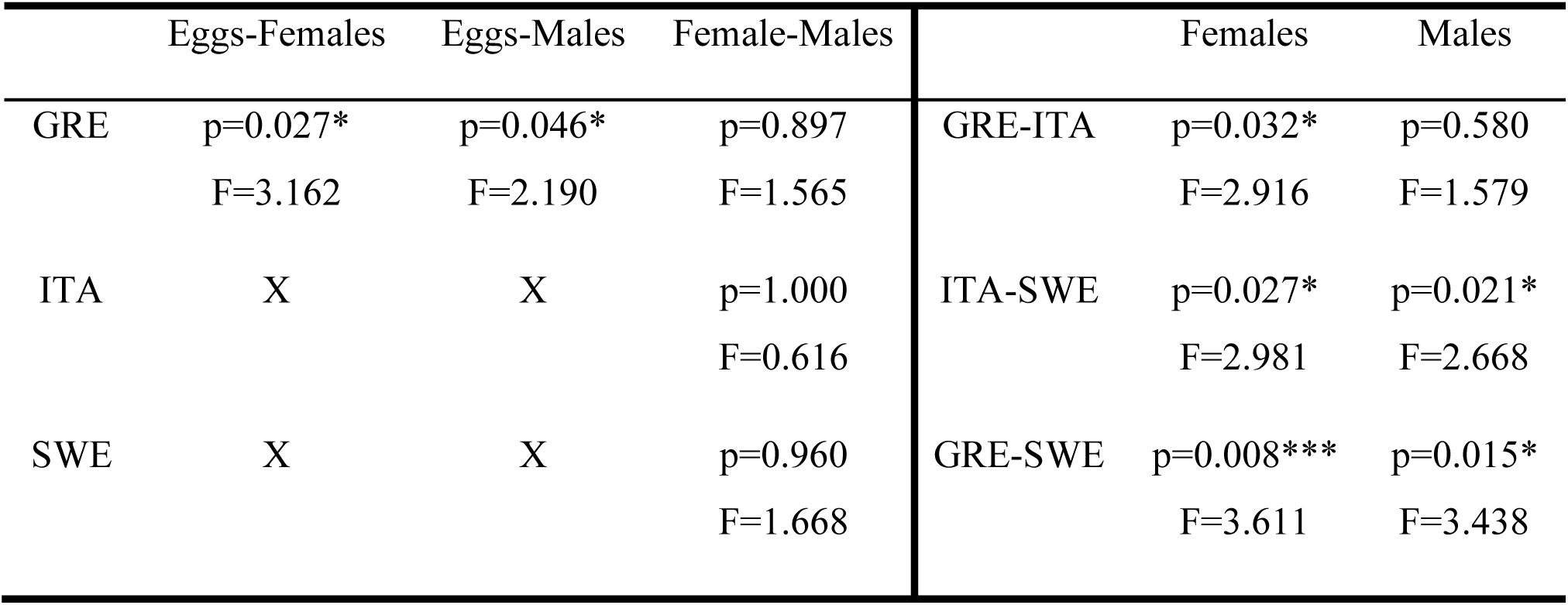
Permutational analysis of variance (PERMANOVA) of eggs, male and female individuals of *Nephrops norvegicus* gut bacterial communities from Greece (GRE), Italy (ITA) and Sweden (SWE). * : p<0.05, ** : p<0.002, *** : p<0.01.

The populations from Italy and Sweden were highly dominated by the same OTU for both females and males, a Photobacterium-related (OTU-0002) and a *Psychrilyobacter*-related (OTU-0001), respectively (Table 1). The females of the Greek population were dominated (17.8%) by the Spirochaetia-related OTU-0004 while the dominant Cyanobiaceae-related OTU-0018 for males prevailed with 8.4% relative abundance (Table S2).

In each population, a different set of OTUs were found to occur in abundances ≥1% concomitantly in both the females and males, i.e. denoted as “most important OTUs” in the study, in each population (Figure 3, Table 3). Three of these, 17 in total, most important OTUs, namely OTU-0001, -0003 and -0013, were found in all three populations. The population from Sweden had 5/10 most important OTUs occurring exclusively in these samples, while the populations from Italy and Greece had only 2/8 and 1/9, respectively. Regression analysis of the most important OTUs abundances vs. their total body weight, showed two cases of statistically significant negative regressions (OTU-0002, -0005) and five cases of positive regressions (OTU-0004, -0010, -0011, - 0013, -0015) (Table 3). There were no “most important” OTUs between the eggs and females or males in the Greek population (Figure S4). In each population, the females:males ratios of abundant OTUs, i.e. relative abundance of ≥1% in each population, which exhibited high (>10) or low (<0.1) values, revealed that various and no overlapping between females and males OTUs contributed differentially to these bacterial communities (Figure 4). Each population had a different set of OTUs with 0.1 10 for all OTUs (Figure S5). The population from Italy had the lowest number of OTUs (8) with females:males ratio >10 and the highest (74) in the population from Sweden. The population from Sweden showed the lowest numbers of OTUs (14) with ratios <0.1 while the highest number of this ratio occurred in the population from Greece (Figure S5).

**Figure 3.**
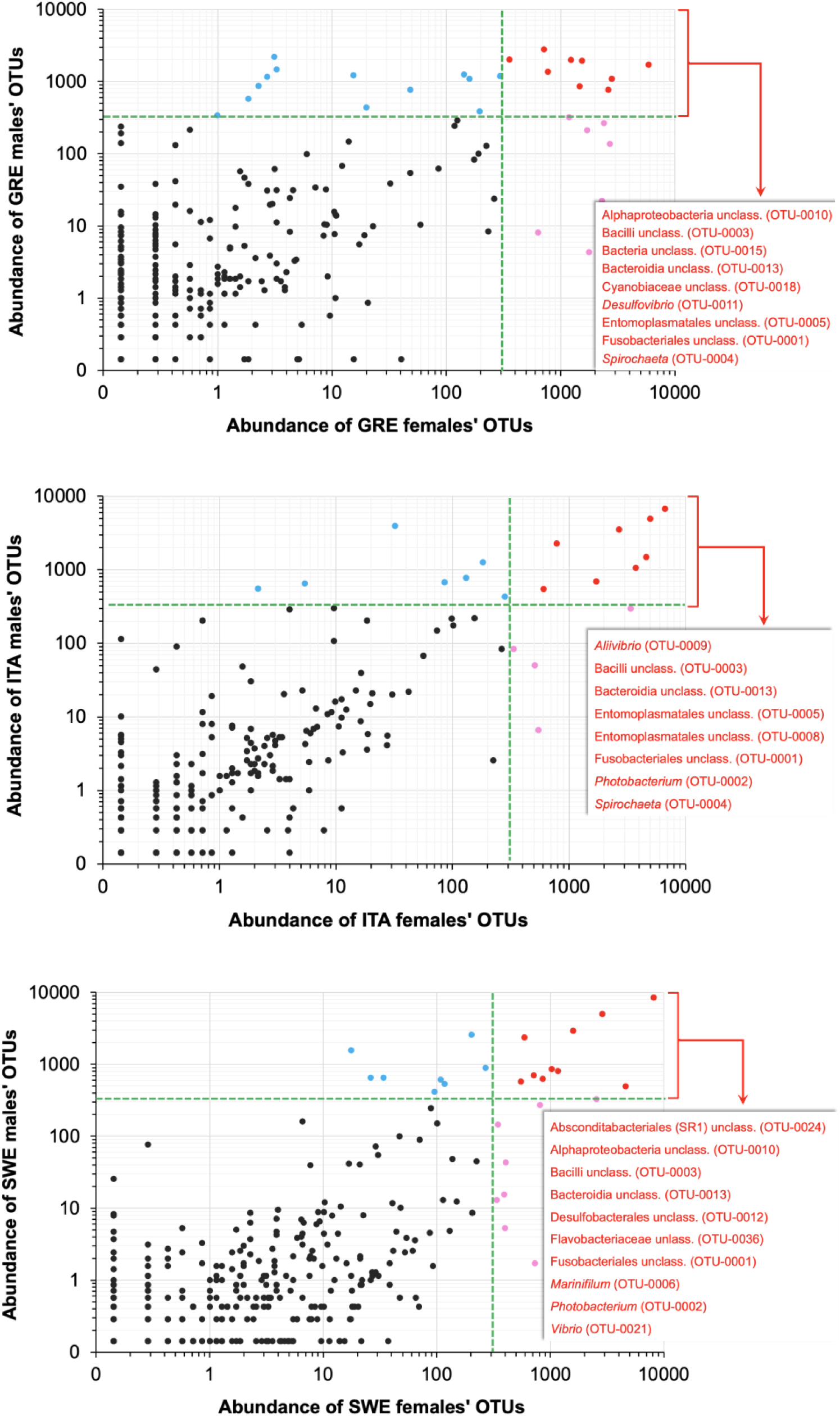
Most important (relative abundance ≥1% in both male and female individuals) bacterial operational taxonomic units (OTUs) of the *Nephrops norvegicus* gut from Greece (GRE), Italy (ITA) and Sweden (SWE). Red dots: bacterial operational taxonomic units (OTUs); blue and pink dots: important OTUs, i.e. ≥1% relative abundance in males and females only, respectively.

**Table 3.**
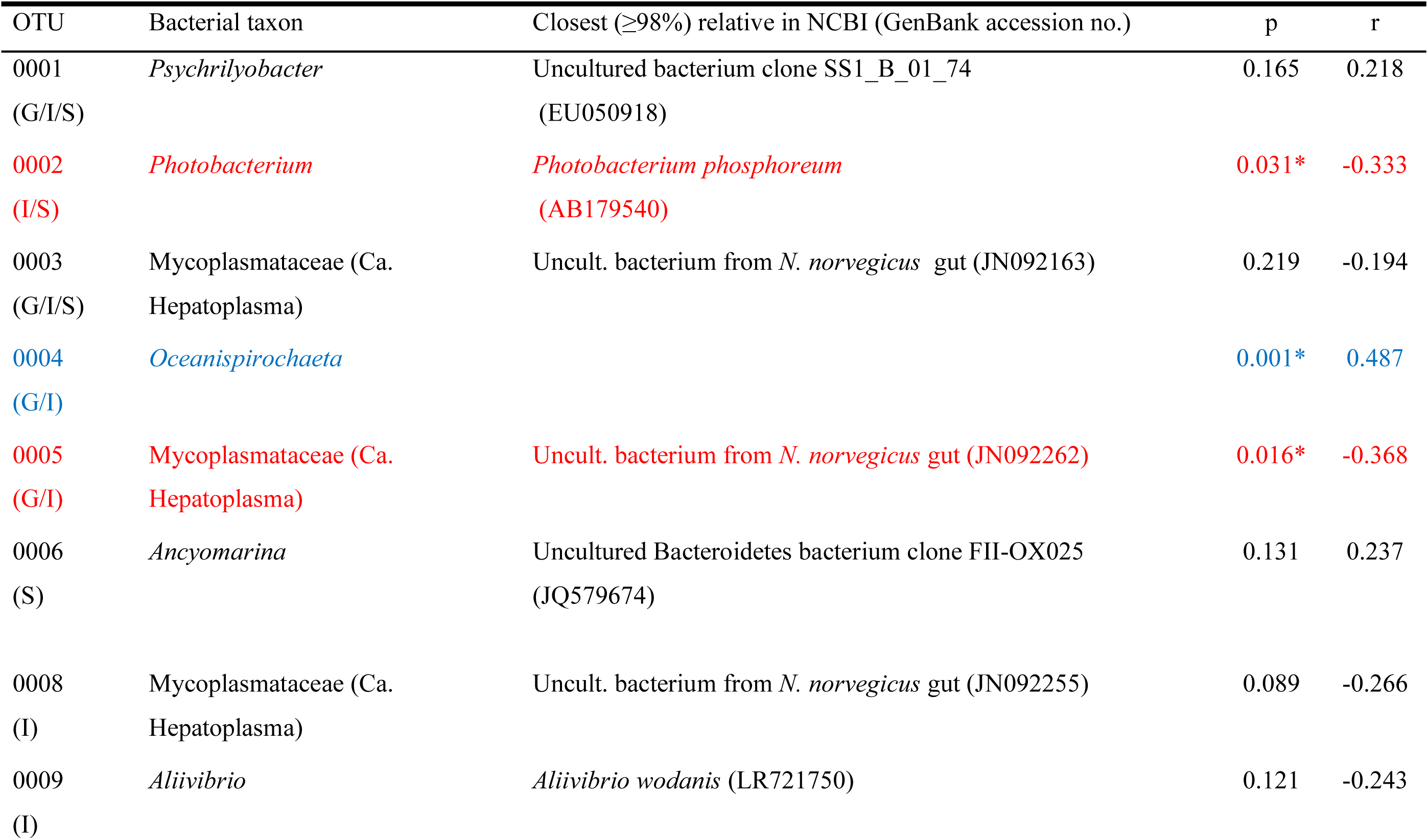
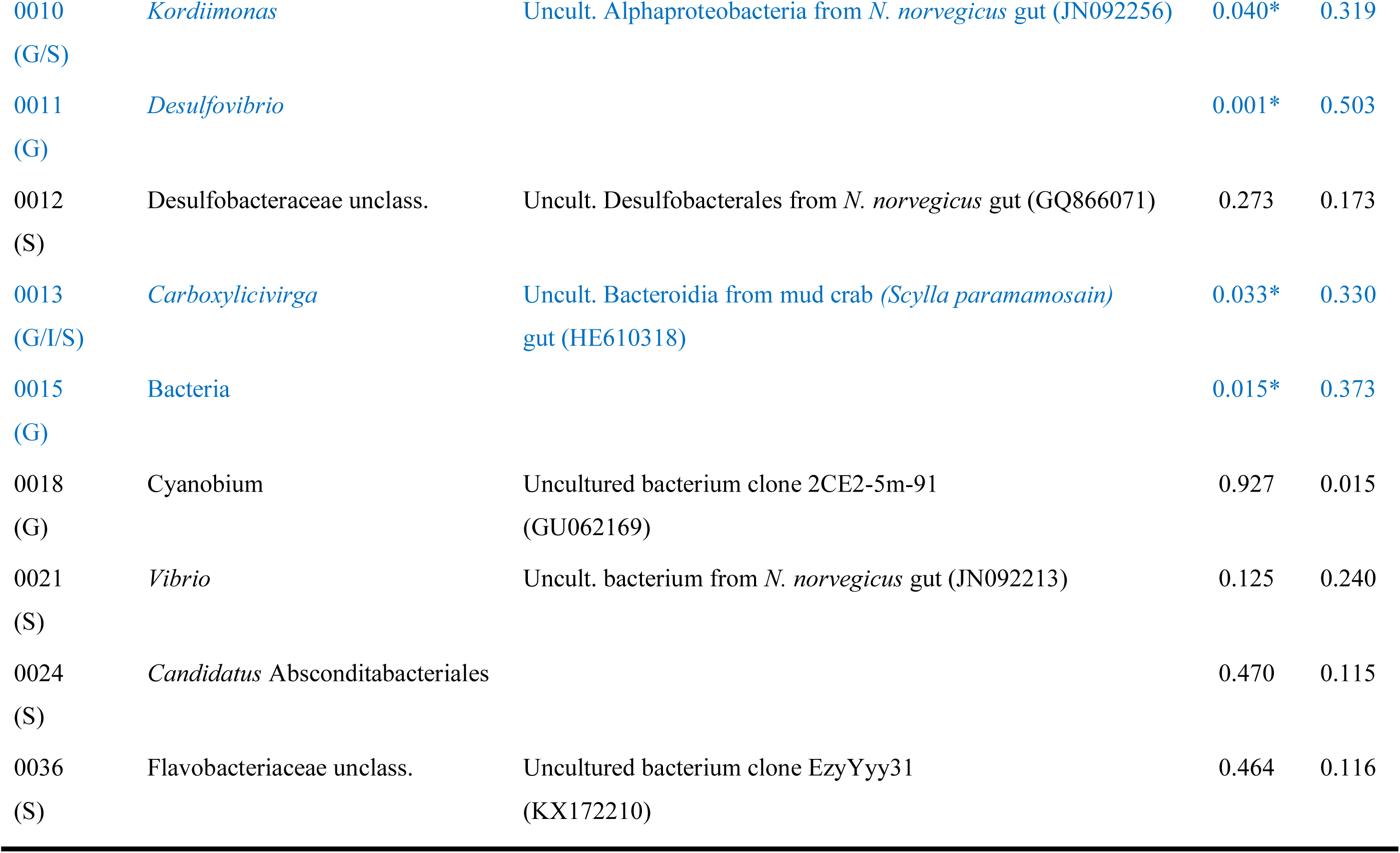
Regression of the total body weight vs. the abundance of each of the most important operational taxonomic units (OTU) from three *Nephrops norvegicus* populations. N=42; negative and positive regressions are in red and blue letters, respectively; *: p<0.05; G: Greece, I: Italy, S: Sweden.

**Figure 4.**
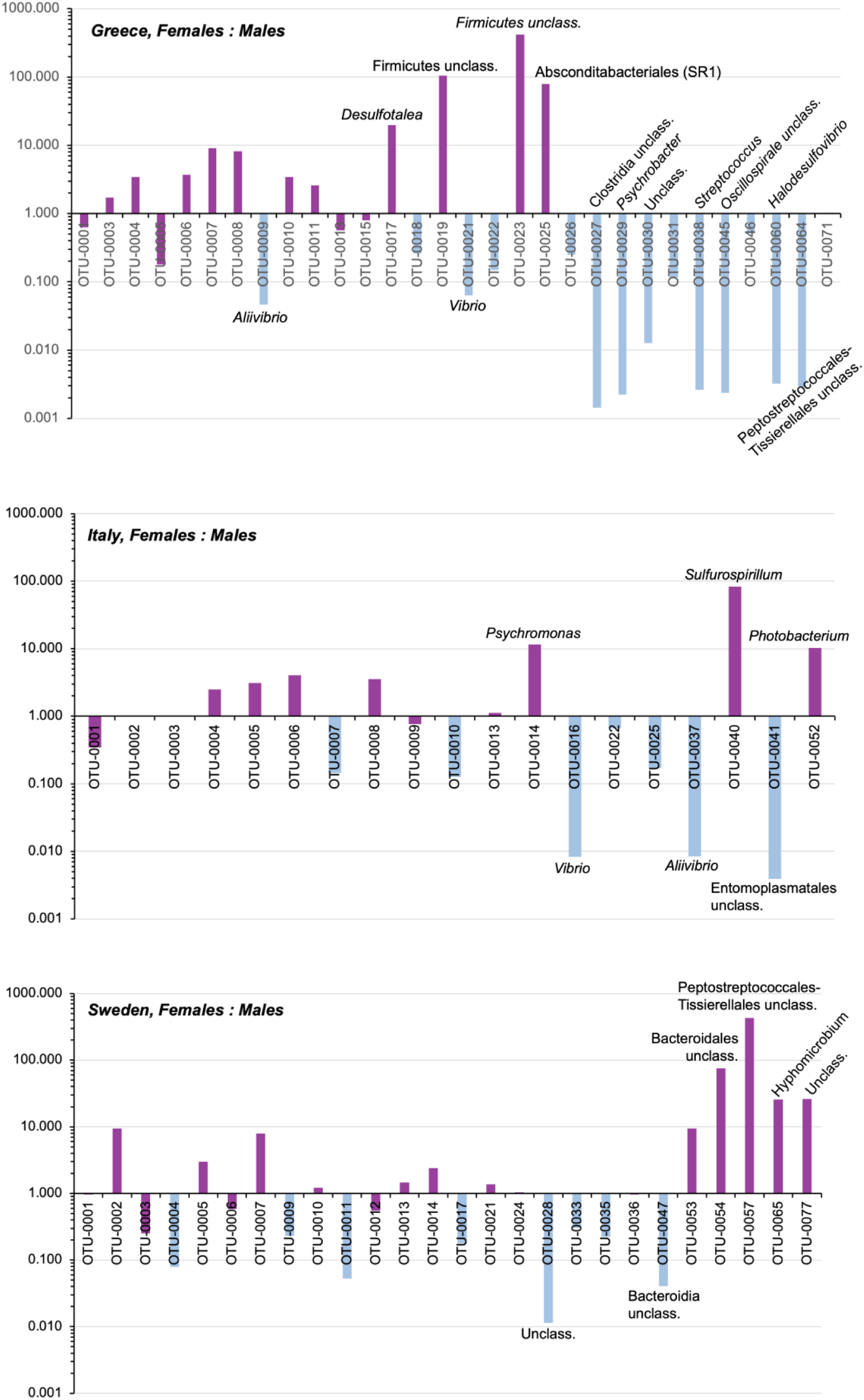
Ratio of females:males bacterial operational taxonomic units (OTUs) with ≥1% relative abundance in the *Nephrops norvegicus* gut from three populations. Only closest relatives of OTUs with 0.1≤ratio≥10 is shown. Blue bars indicate OTUs occurrence only in males.

## 4. DISCUSSION

Our study analyzed the gut bacterial communities in three different *Nephrops norvegicus* populations including both male and female specimens, in order to detect potential differences between geographic regions and sex. As the exact age/developmental stage of the collected animals is difficult to safely be determined, we use the total body weight and carapace length as indicative parameters for growth which can be related to some of the occurred OTUs.

Minor differences were observed between male and female individuals in Sweden and Italy populations while differences observed between male and female gut samples from Greece in both Class and OTU level might be associated with females carrying eggs and thus probably consuming different food sources. Especially for *N. norvegicus* the influence of available feed on gut microbiota has been investigated and confirmed in previous studies (Meziti et al., 2012; Meziti et al., 2010).

Overall dominant OTUs detected in this study imply a more algal-related feeding resource for all populations apart from Italy. More specifically the genus *Psychrilyobacter*, that dominated in the Swedish population, is a Fusobacteriota bacterium that is very commonly detected in association with marine animals such as oysters (Fernandez-Piquer et al., 2012), snails (Aronson et al., 2016), mussels (Santibáñez et al., 2022) and crabs (Zhang et al., 2017) while it also exhibits free-living lifestyles being a part of the rare biosphere (Liu et al., 2023; Yadav et al., 2021). A recent study combining cultivation-dependent and independent methods, revealed some functions from an abalone *Psychrilyobacter* isolate where the specific genus is thriving. Amongst others it was revealed that the isolate can utilize monosaccharides and disaccharides but not polysaccharides, implying that it is possibly involved in later fermentation steps but not in the initial food degradation (Liu et al., 2023). They showed that *Psychrilyobacter* is a versatile fermenter that in collaboration with other bacteria is very important in digesting the algae consumed by the host. However, *N. norvegicus* does not feed on algae although remains of plants have been found in its stomach (Cristo and Cartes, 1998). Previous studies on seasonal changes of the *N. norvegicus* gut microbiome had suggested that temporal changes in water column leading to increased algal material concentration in sediment might enhance the presence of alginolytic communities in *N. norvegicus* gut microbiome (Meziti et al., 2010).

Populations from Greece were characterized by sexual variability leading to the prevalence of a conspicuous Spirochaetaceae representative in females. The genus *Oceanispirochaeta,* observed in this study, contains three fully described species so far, that are obligately anaerobic sediment chemoorganotrophs mainly relying on mono- and disaccharides fermentation (Dubinina et al., 2020; Subhash and Lee, 2017). Although the importance of *Spirochaeta* in the termite gut is well studied (Breznak and Leadbetter, 2006) there is not much information regarding their contribution in marine animals. Recently it has been reported that a spirochaetes associated with a marine sponge mediated terpene production for possible protection of their host against oxidative stress (Waterworth et al., 2024) and has also been speculated to be involved in nitrite metabolism and degradation of complex sugars in the gut of the mudshrimp *Gilvossius tyrrhenus* (formerly *Pestarella tyrrhena*) (Demiri et al., 2009).

The genus *Photobacterium* is commonly detected in *N. norvegicus* and other crustaceans gut microbiome (Foysal, 2023; Jiang et al., 2023; Meziti et al., 2012; Moi et al., 2017). This genus can produce several beneficial compounds such as polyunsaturated fatty acids, lipases, esterases and antimicrobial compounds being also a good candidate for probiotic use (Jiang et al., 2023; Le Doujet et al., 2019). In our study this genus was dominant in the Italian population and important for both Swedish and Italian populations. Its negative correlation with size might imply its prevalence in younger populations. Most importantly, in an older study examining *N. norvegicus* gut microbiota in reared populations receiving different feed, *Photobacterium* was the most abundant genus in mussel fed individuals appearing also as a potential probiotic (Meziti et al., 2012).

Finally, the conspicuous Mycoplasmataceae (ca. Hepatoplasma) representatives were also prevalent in this study similarly to previous studies on *N. norvegicus* as well as on other crustaceans gut microbiome (Meziti et al., 2012). Initially, these Bacteria were detected in the midgut glands (hepatopancreas) of the terrestrial isopod *Porcellio scaber* (Wang et al., 2004) and it has been shown to benefit their hosts under low-nutrient conditions (Fraune and Zimmer, 2008). Today it seems that this bacterial taxon seems to be frequently abundant in various decapods (Foysal, 2023), such as the hepatopancreas of the velvet crab *Necora puber* (Martin et al., 2024), in the gut of deep-sea amphipods (Cheng et al., 2019) juvenile (Sun et al., 2020) and adult individuals (An et al., 2024) of the mitten crab *Eriocher sinensis*, juvenile Caribbean spiny lobsters *Panulirus argus* (Zamora-Briseño et al., 2020), mud crab *Scylla paramamosain* (Jiang et al., 2023) and the vent shrimp *Rimicaris exoculata* (Aubé et al., 2022). Genome sequencing of ca. Hepatoplasma (Collingro et al., 2015) as well as Metagenome Assembled Genomes (MAGs) sequencing (Aubé et al., 2022), showed typical Mycoplasmataceae reduced genome size with the majority of energy producing pathways missing with the exception of glycolysis. Similarly, the majority of nucleotide and amino acid biosynthesis pathways are not present in the genomes suggesting that ca. Hepatoplasma mainly relies on its host or syntrophic bacteria for its growth (Aubé et al., 2022).

Overall the majority of the dominant and ‘important’ OTUs in our dataset belonged to groups and genera that had been previously detected in *N. norvegicus* or other Crustaceans gut microbiome studies (Meziti et al., 2012; Meziti et al., 2010); (Aubé et al., 2022; Jiang et al., 2023). Most importantly, the closest relatives detected, for the majority of the ‘important OTUs’, were phylotypes from previous *N. norvegicus* gut microbiome studies mainly performed in Greece more than one decade ago implying the presence of a core gut *N. norvegicus* microbiome regardless of geographical boundaries. Similar findings were reported for the mud crab *Scylla paramosain,* where ca. Hepatoplasma, *Vibrio, Photobacterium, Carboxylicivirga* were identified as core gut microbiota genera from different coastal regions in southern China (Jiang et al., 2023).

To date, no known interaction between the *N. norvegicus* sex and its gut microbiota exists. However, it is known that some vertebrate steroids (e.g. estradiol, progesterone, testosterone) influence and are dependent on the developmental stage and reproduction of the *H. americanus* and *N. norvegicus* lobsters; these steroids are most likely species-specific (Burns et al., 1984; Chang, 1997; Fairs et al., 1989). Bacteria seem to have a role in steroid hormone levels in mammals. For example, the activity of the β-glucuronidase produced by gut bacteria, like *Bacteroides* and *Clostridium,* regulate the levels of active estrogen by breaking glycosidic bonds between glucuronic acid and estrogen (Cotton et al., 2023). β-glucuronidases are widespread along several bacterial taxa (Lombard et al., 2013; Wardman et al., 2022), some of which are among the most abundant found in this study (e.g. Clostridia, Sphingomonadaceae, Spirochaetia). Bacteria seem to be also involved in the degradation or the production/reactivation of testosterone while gut microbiota changes can be accompanied by testosterone levels (Cotton et al., 2023). Other invertebrates, such as cephalopods, have an accessory nidamental gland which is known to harbour specific microbiota for assisting the defense of their females against pathogens and fouling organisms (Vijayan et al., 2024). In addition, our current dataset cannot provide any potential metabolic features of the investigated microbial communities as among the most important OTUs several cannot be securely affiliated to any of the known bacterial taxa (see Table S2). These yet-to-be cultivated Bacteria, which hinder the full characterization of their ecological demands, perhaps refrain us from having a more complete picture for the life cycle of this animal but also for its commercial cultivation.

## AKNOWLEDGEMENTS

We thank Barbara Bland and Jan-Erik Johansson at the Swedish University of Agricultural Sciences for the collection of the North Sea samples onboard the R/V “SVEA”, Danilo Scannella (IRBIM CNR) for the collection of the Strait of Sicily samples and Vasiliki Kousteni (Fisheries Research Institute, Hellenic Agricultural Organization - Demeter, Kavala, Greece) for the collection of the North Aegean Sea samples.

