## Supplementary material for "Sex-related gut microbiota in three geographically separated Norway lobster (*Nephrops norvegicus*) populations"

**Table S1**. Morphometric data of the analysed *Nephrops norvegicus* specimens.

| | Carapace length (mm) | Total Weight (g) | Sex |
|---|---|---|---|
| | 49.91 | 80.7 | ♀ |
| | 36.83 | 37.8 | ♀ |
| | 46.29 | 78.8 | ♀ |
| | 44.3 | 67.3 | ♀ |
| | 44.9 | 74.5 | ♀ |
| | 45.51 | 64.4 | ♀ |
| Greece | 54.15 | 114.3 | ♀ |
| | 51.41 | 87.9 | ♂ |
| | 38.95 | 37.9 | ♂ |
| | 44.79 | 69.2 | ♂ |
| | 36.25 | 37.7 | ♂ |
| | 51.79 | 104.7 | ♂ |
| | 53.4 | 124.7 | ♂ |
| | 50.33 | 77.1 | ♂ |
| | 37 | 35.3 | ♀ |
| | 33 | 32.5 | ♀ |
| | 36 | 33.7 | ♀ |
| | 33 | 29.6 | ♀ |
| | 36 | 37.5 | ♀ |
| | 35 | 28.1 | ♀ |
| Italy | 34 | 30.2 | ♀ |
| | 37 | 39.9 | ♂ |
| | 34 | 30.7 | ♂ |
| | 37 | 35.7 | ♂ |
| | 37 | 40.6 | ♂ |
| | 39 | 42.2 | ♂ |
| | 37 | 28.9 | ♂ |
| | 37 | 36.4 | ♂ |
| | 45 | 65.0 | ♀ |
| | 45 | 66.0 | ♀ |
| | 45 | 63.0 | ♀ |
| | 45 | 67.0 | ♀ |
| | 45 | 65.0 | ♀ |
| | 45 | 63.0 | ♀ |
| Sweden | 45 | 63.0 | ♀ |
| | 45 | 66.0 | ♂ |
| | 45 | 68.0 | ♂ |
| | 45 | 67.0 | ♂ |
| | 45 | 69.0 | ♂ |
| | 45 | 68.0 | ♂ |
| | 45 | 69.0 | ♂ |
| | 45 | 68.0 | ♂ |

**Table S2**. Midgut bacterial operational taxonomic units (OTU) comprising ≥80% cumulative relative abundance in *Nephrops norvegicus* populations.

| OTU | Taxon | Eggs | OTU | Taxon | Females | OTU | Taxon | Males |
|---|---|---|---|---|---|---|---|---|
| | | GREECE | | | GREECE | | | GREECE |
| 0020 | *Cutibacterium* | 18.5% | 0004 | *Spirochaeta*-2 | 17.8% | 0018 | Cyanobiaceae unclass. | 8.4% |
| 0032 | *Methylobacterium-Methylorubrum* | 7.1% | 0011 | *Desulfovibrio* | 8.5% | 0027 | Clostridia unclass. | 6.6% |
| 0039 | *Sphingomonas* | 7.1% | 0017 | *Desulfotalea* | 8.2% | 0005 | Entomoplasmatales unclass. | 6.0% |
| 0034 | Comamonadaceae unclass. | 7.0% | 0010 | Alphaproteobacteria unclass. | 7.9% | 0001 | Fusobacteriales unclass. | 5.9% |
| 0044 | *Fusobacterium* | 4.7% | 0007 | Bacilli unclass. | 7.2% | 0015 | Bacteria unclass. | 5.8% |
| 0051 | *Acinetobacter* | 4.1% | 0019 | Firmicutes unclass. | 7.0% | 0004 | *Spirochaeta*-2 | 5.1% |
| 0043 | Neisseriaceae unclass. | 4.0% | 0023 | Firmicutes unclass. | 5.4% | 0029 | *Psychrobacter* | 4.4% |
| 0055 | Corynebacteriales unclass. | 4.0% | 0008 | Entomoplasmatales unclass. | 5.2% | 0013 | Bacteroidia unclass. | 4.1% |
| 0042 | *Chryseobacterium* | 3.9% | 0015 | Bacteria unclass. | 4.6% | 0031 | Bacteroidia unclass. | 3.7% |
| 0048 | *Corynebacterium* | 3.5% | 0003 | Bacilli unclass. | 4.4% | 0030 | Bacteria unclass. | 3.7% |
| 0050 | *Staphylococcus* | 3.2% | 0001 | Fusobacteriales unclass. | 3.7% | 0026 | *Synechococcus* | 3.6% |
| 0056 | *Streptococcus* | 3.0% | 0006 | *Marinifilum* | 3.6% | 0045 | Oscillospirales unclass. | 3.5% |
| 0063 | Rhizobiaceae unclass. | 3.0% | | | | 0011 | *Desulfovibrio* | 3.3% |
| 0078 | *Mesorhizobium* | 2.2% | | | | 0022 | Arcobacteraceae unclass. | 3.3% |
| 0067 | *Pseudomonas* | 2.1% | | | | 0038 | *Streptococcus* | 2.6% |
| 0068 | *Rothia* | 1.9% | | | | 0003 | Bacilli unclass. | 2.6% |
| 0062 | Micrococcaceae unclass. | 1.7% | | | | 0010 | Alphaproteobacteria unclass. | 2.3% |
| | | | | | | 0021 | *Vibrio* | 2.3% |

| ***OTU*** | ***Taxon*** | ***Females*** | ***OTU*** | ***Taxon*** | ***Males*** |
|---|---|---|---|---|---|
| | | ITALY | | | ITALY |
| 0002 | *Photobacterium* | 20.2% | 0002 | *Photobacterium* | 20.3% |
| 0003 | Bacilli unclass. | 15.1% | 0003 | Bacilli unclass. | 14.8% |
| 0005 | Entomoplasmatales unclass. | 13.9% | 0016 | *Vibrio* | 11.8% |
| 0008 | Entomoplasmatales unclass. | 11.3% | 0009 | *Aliivibrio* | 10.6% |
| 0014 | *Psychromonas* | 10.3% | 0001 | Fusobacteriales unclass. | 6.8% |
| 0009 | *Aliivibrio* | 8.1% | 0005 | Entomoplasmatales unclass. | 4.5% |
| 0004 | *Spirochaeta*-2 | 5.2% | 0007 | Bacilli unclass. | 3.8% |
| 0001 | Fusobacteriales unclass. | 2.4% | 0008 | Entomoplasmatales unclass. | 3.2% |
| 0013 | Bacteroidia unclass. | 1.8% | 0025 | Bacilli unclass. | 2.3% |
| 0040 | *Sulfurospirillum* | 1.7% | 0004 | *Spirochaeta*-2 | 2.1% |
| 0052 | *Photobacterium* | 1.5% | 0010 | Alphaproteobacteria unclass. | 2.0% |
| 0006 | *Marinifilum* | 1.0% | 0037 | *Aliivibrio* | 2.0% |
| 0022 | Arcobacteraceae unclass. | 0.9% | 0041 | Entomoplasmatales type-III | 1.7% |
| 0011 | *Desulfovibrio* | 0.8% | 0013 | Bacteroidia unclass. | 1.6% |
| 0089 | *Vibrio* | 0.7% | 0022 | Arcobacteraceae unclass. | 1.3% |
| 0007 | Bacilli unclass. | 0.6% | 0049 | *Shewanella* | 0.9% |
| 0017 | *Desulfotalea* | 0.5% | 0014 | *Psychromonas* | 0.9% |

| | | | | | |
|---|---|---|---|---|---|
| 0025 | Bacilli unclass. | 0.4% | 0058 | *Aliivibrio* | 0.9% |
| 0035 | Desulfocapsaceae unclass. | 0.3% | 0017 | *Desulfotalea* | 0.7% |
| 0021 | *Vibrio* | 0.3% | 0021 | *Vibrio* | 0.7% |

| ***OTU*** | ***Taxon*** | ***Females*** SWEEDEN | ***OTU*** | ***Taxon*** | ***Males*** SWEEDEN |
|---|---|---|---|---|---|
| 0001 | Fusobacteriales unclass. | 24.5% | 0001 | Fusobacteriales unclass. | 25.5% |
| 0002 | *Photobacterium* | 14.0% | 0006 | *Marinifilum* | 15.0% |
| 0006 | *Marinifilum* | 8.7% | 0012 | Desulfobacterales unclass. | 8.8% |
| 0007 | Bacilli unclass. | 7.7% | 0004 | *Spirochaeta*-2 | 7.8% |
| 0012 | Desulfobacterales unclass. | 4.8% | 0003 | Bacilli unclass. | 7.1% |
| 0013 | Bacteroidia unclass. | 3.5% | 0028 | Bacteria unclass. | 4.7% |
| 0010 | Alphaproteobacteria unclass. | 3.1% | 0033 | Bacteroidia unclass. | 2.7% |
| 0021 | *Vibrio* | 2.6% | 0010 | Alphaproteobacteria unclass. | 2.6% |
| 0005 | Entomoplasmatales unclass. | 2.4% | 0013 | Bacteroidia unclass. | 2.4% |
| 0057 | Peptostreptococcales-Tissierellales unclass. | 2.2% | 0024 | *Vibrio* | 2.1% |
| 0024 | *Vibrio* | 2.2% | 0011 | Desulfovibrio | 2.0% |
| 0003 | Bacilli unclass. | 1.8% | 0047 | Bacteroidia unclass. | 1.9% |
| 0036 | Flavobacteriaceae unclass. | 1.7% | 0021 | *Vibrio* | 1.9% |
| 0053 | Terasakiellaceae unclass. | 1.2% | 0017 | *Desulfotalea* | 1.8% |
| 0054 | Bacteroidales unclass. | 1.2% | 0036 | Flavobacteriaceae unclass. | 1.7% |

| | | | | | | |
|---|---|---|---|---|---|---|
| | 0065 | *Hyphomicrobium* | 1.2% | 0035 | Desulfocapsaceae unclass. | 1.6% |
| | 0014 | *Psychromonas* | 1.0% | 0002 | *Photobacterium* | 1.5% |
| | 0077 | Bacteria unclass. | 1.0% | 0009 | *Aliivibrio* | 1.3% |
| | 0033 | Bacteroidia unclass. | 0.8% | 0007 | Bacilli unclass. | 1.0% |
| | 0074 | Actinomarinales unclass. | 0.7% | 0005 | Entomoplasmatales unclass. | 0.8% |

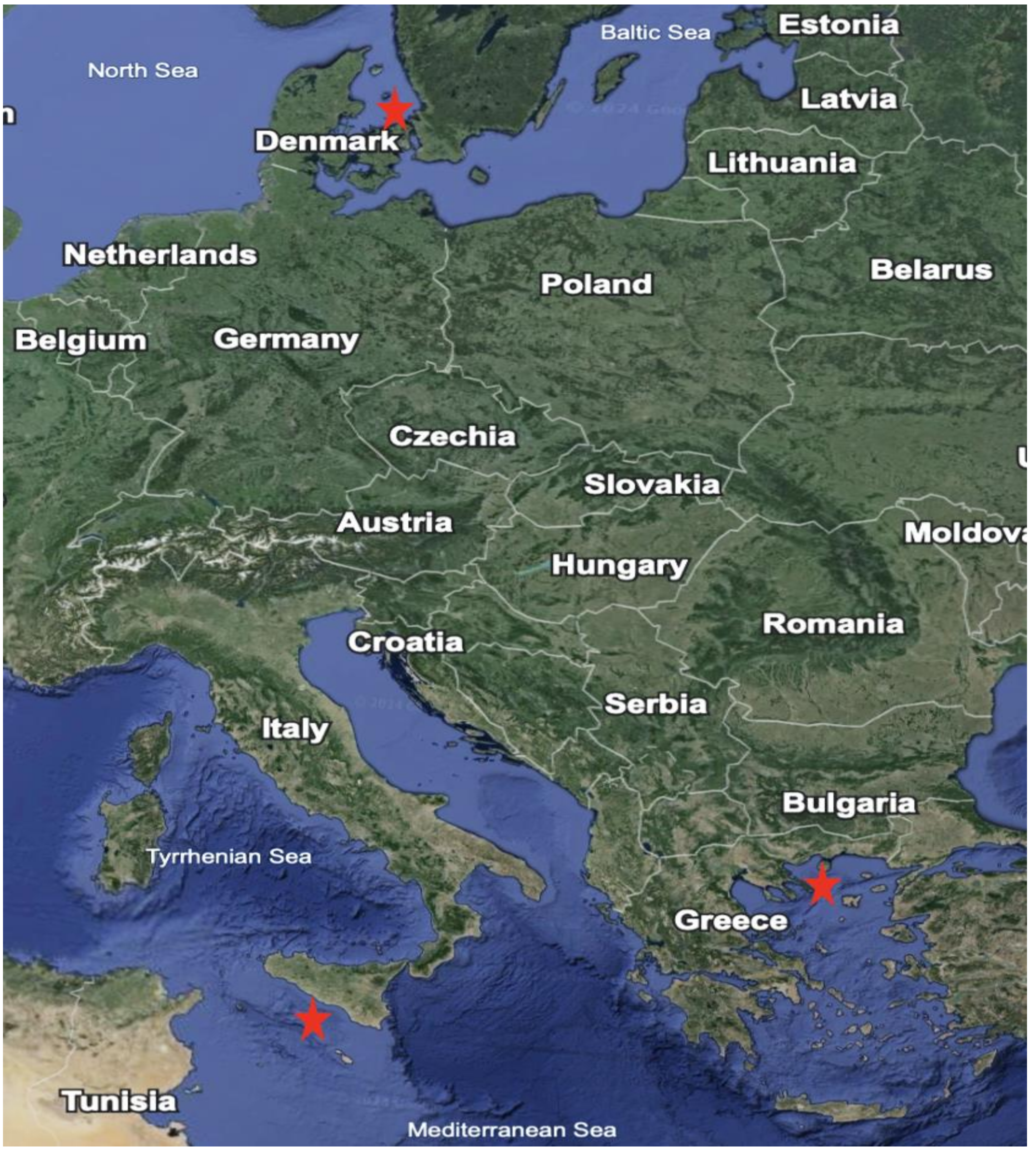


**Figure S1**. Map of sampling points (red stars).

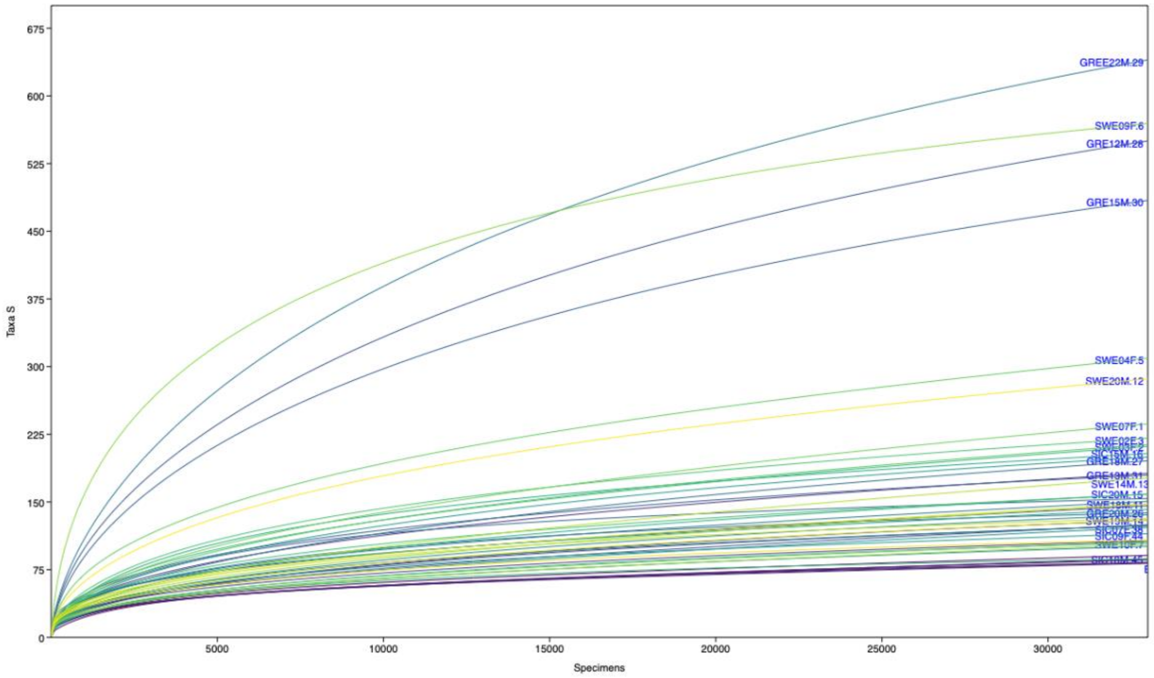


**Figure S2**. Rarefaction curves of all midgut samples from three *Nephrops norvegicus* populations.

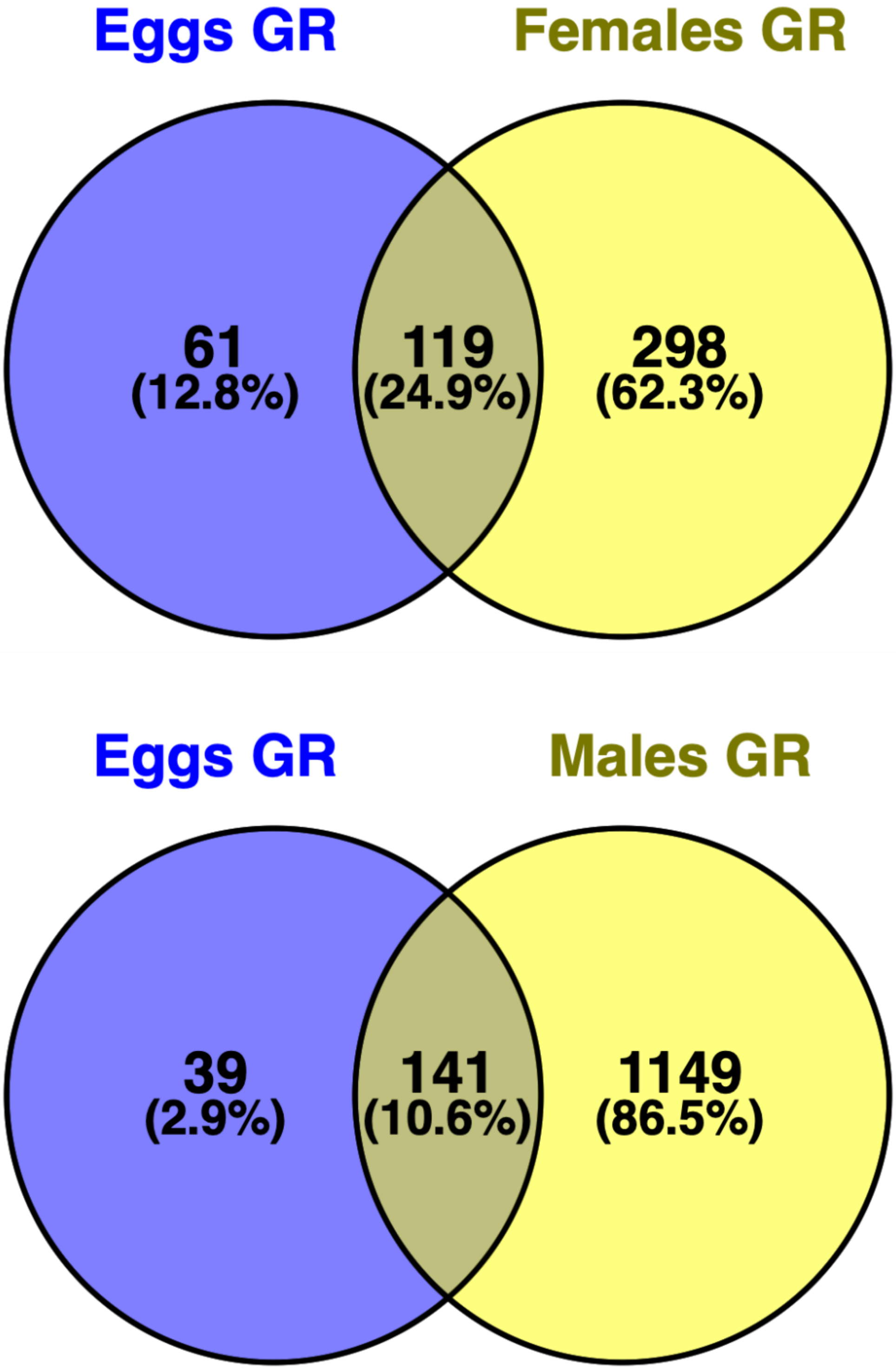


**Figure S3**. Shared bacterial operational taxonomic units between eggs and female/male specimens in a *Nephrops norvegicus* populations from Greece.

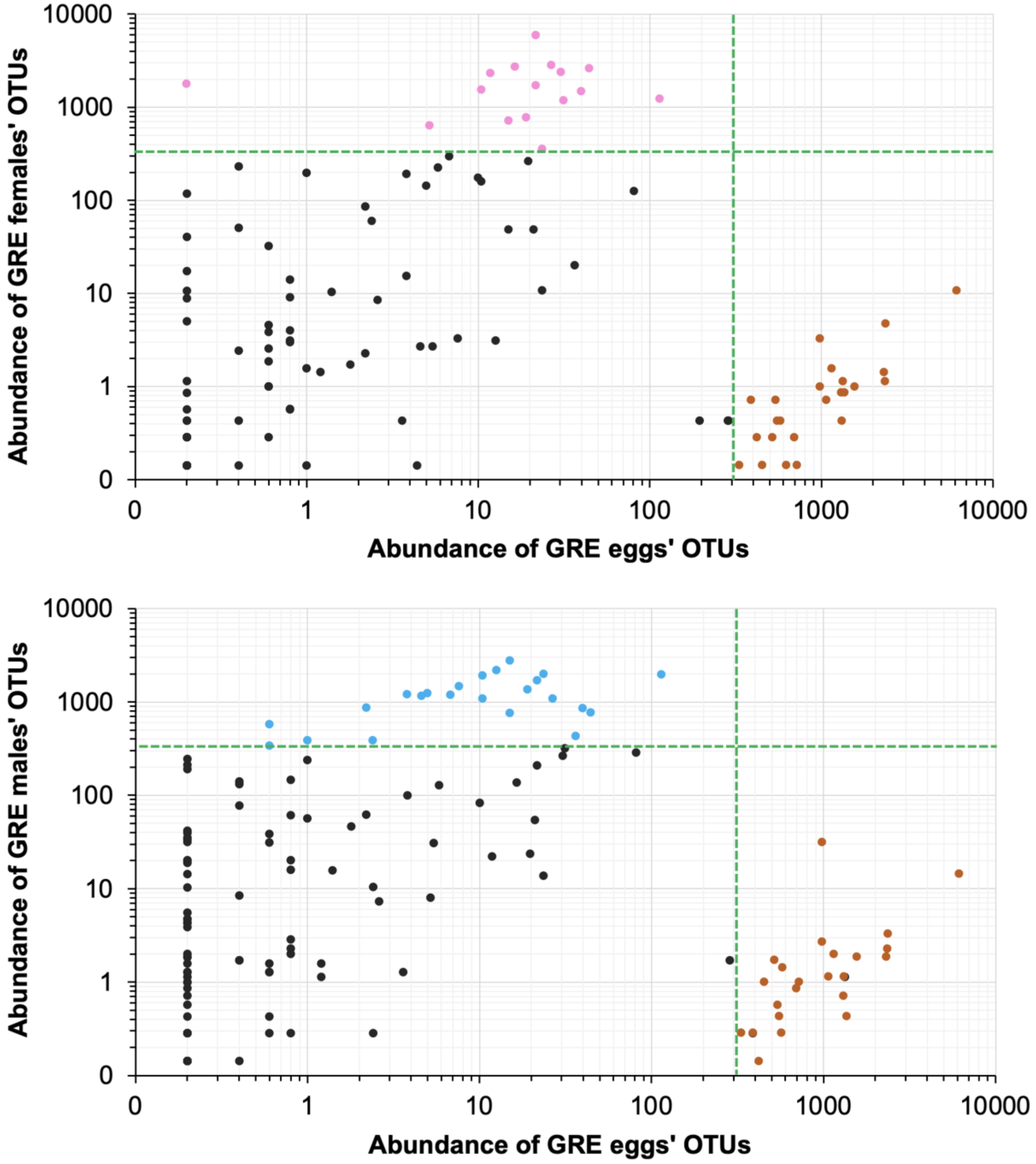


**Figure S4**. Important (relative abundance ≥1%) bacterial operational taxonomic units (OTUs) in Norway lobster's gut from Greece (GRE) between eggs and adults. Brown dots indicate important OTUs in eggs only. Blue and pink dots indicate important OTUS in males and females only, respectively.

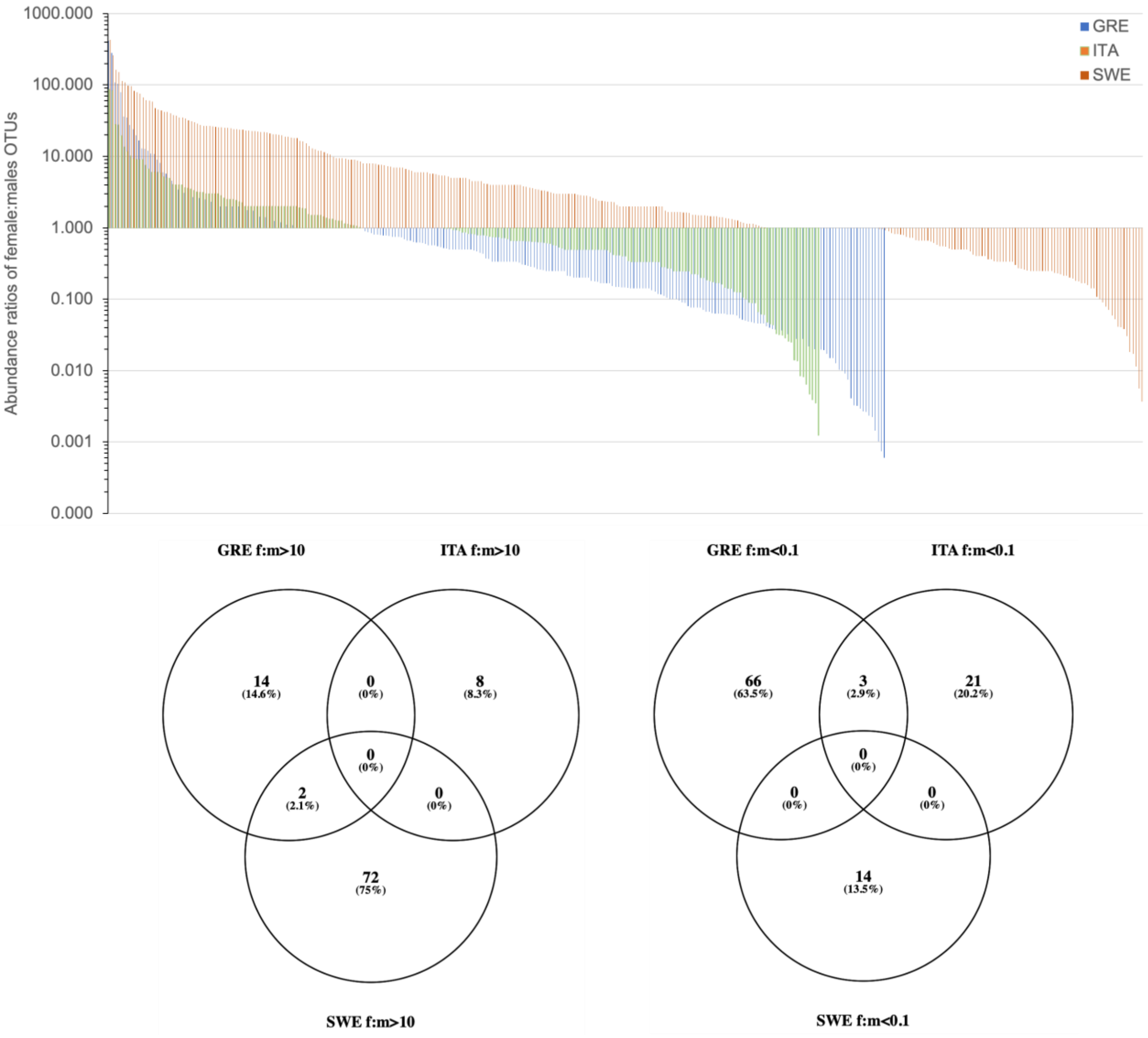


**Figure S5**. The females:males ratio of bacterial operational taxonomic units (OTUs) abundance in three *Nephrops norvegicus* populations.
